# Show the model – A Harrell plot combines a forest plot of effects and uncertainty with a dot plot of the data

**DOI:** 10.1101/458182

**Authors:** Jeffrey A. Walker

## Abstract

Several high profile editorials and articles have advocated replacing the ubiquitous, but not especially informative, bar plot with dot plots or box plots which show the data or some summary of the data distribution. I argue that the biggest deficiency of bar plots (and other mean-and-error plots) is that they fail to directly communicate modeled effects and uncertainty, and dot and box plots do not address this deficiency. A Harrell plot combines a forest plot of modeled treatment effects and uncertainty, a dot plot of raw data, and a box plot of the distribution of the raw data into a single, two-part graph. A Harrell plot encourages best practices such as communication of the distribution of the data and a focus on effect size and uncertainty, while discouraging poor practices such as hiding distributions and focusing on *p*-values.

## Introduction

Recommended best practices for the reporting of statistical results include 1) showing the raw data and/or distribution of data in plots (Drummond and Vowler 2011; Weissgerber et al. 2015; Spitzer et al. 2014; Krzywinski and Altman 2014; Harrell 2014; Weissgerber et al. 2016) and focusing on 2) effect size and (3) uncertainty in effect estimates instead of *p*-values of null hypothesis tests (Nakagawa and Cuthill 2007; Yoccoz 1991; Johnson 1999; Curran-Everett, Taylor, and Kafadar 1998). By contrast, standard practice throughout experimental biology includes the reporting of ANOVA results in tables and treatment means and standard errors of the mean in plots. At best, ANOVA tables poorly communicate effect size and uncertainty. Effects and uncertainty can be inferred from plots of treatment means and standard errors only indirectly.

Here, I introduce the Harrell plot, a tool to communicate statistical results from experiments, or any analysis with categorical independent variables (“ANOVA-like” linear models, including linear mixed models and generalized linear models). A Harrell plot combines 1) a dot plot to show individual values, 2) either a box plot or a mean-and-CI plot to show a summary of the distribution of the response within treatment groups, and 3) a forest plot of effect estimates and confidence intervals to show modeled effect sizes and uncertainty. The combination of the effects in the top part and distribution in the bottom part of a Harrell plot was inspired by Fig. 1.1 of Harrell (2014). The Harrell plot is implemented as an online

**HarrellPlot Shiny app** for users with no or limited R experience, including undergraduate biology majors. An R package for users with some R experience can be downloaded using install_github(“middleprofessor/harrellplot”). The R markdown file to reproduce all figures is available as a supplement.

## Effect size and uncertainty

By effect, or effect size, I mean the magnitude and direction of the difference in response to some treatment, or some combination of treatments. If the mean critical thermal minimum is 5.1 ° C in the control group of flies and 5.8 ° C in the treated group, then the effect is 5.8°C – 5.1C = +0.7°C. The non-intercept coefficients of a linear model are effects. Differences between conditional or marginal means from a statistical model are effects. A confidence interval of the effect is a measure of the uncertainty in the estimate. A 95% confidence interval of the effect has a 95% probability (in the sense of long-run frequency) of containing the true effect. This probability is a property of the population of intervals that could be computed using the same sampling and measuring procedure. It is not correct, without further assumptions, to state that there is a 95% probability that the true effect lies within the interval. However, if we have only weak prior beliefs about the possible values of the effect, then it is valid, though possibly misleading, to state that there is an approximately 95% probability that the true effect lies in the interval (Greenland and Poole 2013; Gelman 2013). Perhaps a more useful interpretation is that the interval contains the range of effects that are consistent with the data, in the sense that a *t*-test would not reject the null hypothesis of a difference between the estimate and any value within the interval (this interpretation does not imply anything about the true value).

While many experiments in biology are conducted with a proximate goal of discovering or confirming that an effect exists (that is, the effect is something other than zero), the ultimate goal of a research program should be to understand the biological (including clinical, behavioral, ecological or evolutionary) consequences of effects (Yoccoz 1991; Curran-Everett, Taylor, and Kafadar 1998; Nakagawa and Cuthill 2007; Batterham and Hopkins 2006). These consequences are functions of effect magnitude and direction and, consequently, estimates of effect size and uncertainty are tools for these ultimate goals. By contrast, the value of hypothesis testing and *p*-values is limited to the more proximate goal of effect presence (and, because frequentist hypothesis testing only provides evidence against a null, it only provides indirect evidence for an effect). Importantly, the discovery that an effect exists requires more than a *p*-value, including both replicate experiments and modified experiments that “probe their experimental systems in multiple, independent ways” (Vaux 2012; see also Munafò and Davey Smith 2018). Probing is standard in much of cell and molecular biology (but see Kaelin Jr 2017). Replications are uncommon throughout most of experimental biology. It probably cannot be emphasized enough that finding statistically significant *p*-values is a very low-bar in experimental biology. All components of complex biological systems are causally connected and perturbing any feature of a system will have some effect on everything, however small (although true null effects may occur in experimental systems with a minimal number of components). While *p*-values are a useful, if limited, tool in a researchers toolbox, Type I errors will generally not exist in complex systems and significance testing (comparing a *p*-value to a Type I error rate, the binning of results into “significant” or “non-significant”, using asterisks to indicate “levels of significance”) should be avoided. Instead, researchers should be concerned with sign and magnitude errors (Gelman and Carlin 2014) and with conditional dependencies.

Confidence intervals of effects can be (and are most often) used to infer “statistical significance”. Statisticians have long advocated for the far more valuable use of a confidence interval as a tool to infer the sensitivity of an interpretation or conclusion to the data (Tukey 1991). Again, a confidence interval of an effect gives the range of parameter values that are consistent with the data (Amrhein, Korner-Nievergelt, and Roth 2017). Consequently, as evidence for a theory in academic biology or a decision in applied biology, the whole range of values within a confidence interval, and not just the point estimate, should be consistent with an interpretation or conclusion, otherwise the data are ambiguous (or “inconclusive”, but this might suggest that the results from a single study could ever be “conclusive”).

### Motivating data – Effects of increased temperature and CO_2_ on larval sea urchin growth and metabolism

The Harrell plot was motivated by the reported results of an experiment measuring the effect of increased temperature and CO_2_ on metabolic rate (among other outcomes) in the larvae of the urchin *Strongylocentrotus purpuratus* (Padilla-Gamiño et al. 2013). The factorial design included two levels of *Temp* (13°C and 18°C) and two levels of CO_2_ (400 μATM and 1100 μATM) with *n* = 6 replicates in each treatment combination. The motivating part of the study was the *p*-value of the interaction effect, which was 0.079 and raised the question “what is the best practice for interpreting results like this?” The data plotted in Figures 1 and 2 here are simulated to closely match the results reported in Fig. 2 of Padilla-Gamiño et al. (2013)(the original data are not available in a public archive). Here, I’ve recoded the *Temp* factor levels to “Temp-” and “Temp and the *CO*_2_ facctor levels to”CO2-" and “CO2+”.

**Figure 1:**
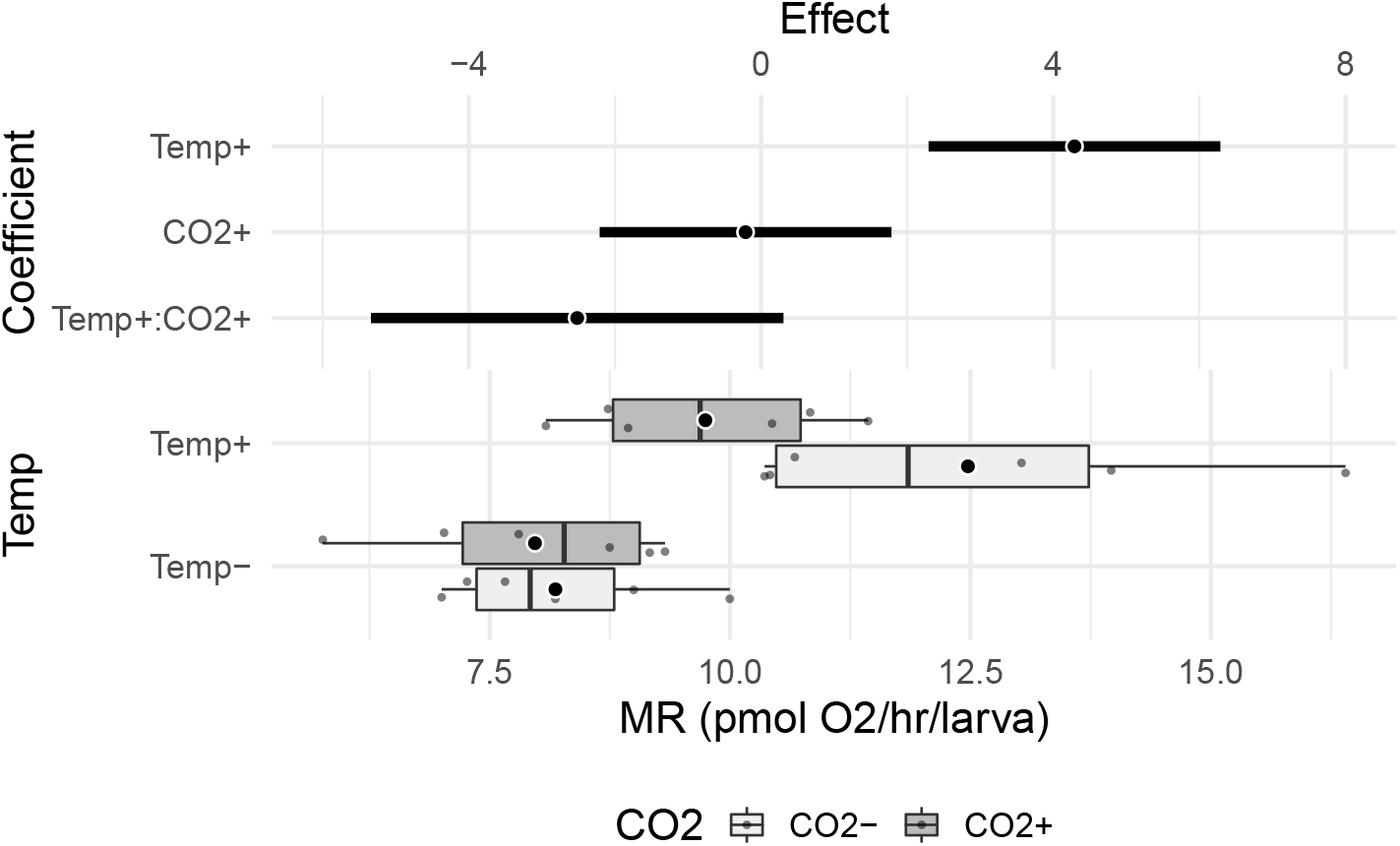
Harrell plot of the simulated urchin data. The plot has two parts. The lower part is a superimposed dot plot and box plot. The large circle for each treatment combination is the conditional mean. The upper part shows the effects and effect uncertainty. Here the effects are parameter estimates (coefficients) of the model and 95% confidence intervals of these estimates. Key: Temp-= 13°C, Temp+ = 18°C, CO2-= 400 μATM, CO2+ = 1100 μATM.

**Figure 2:**
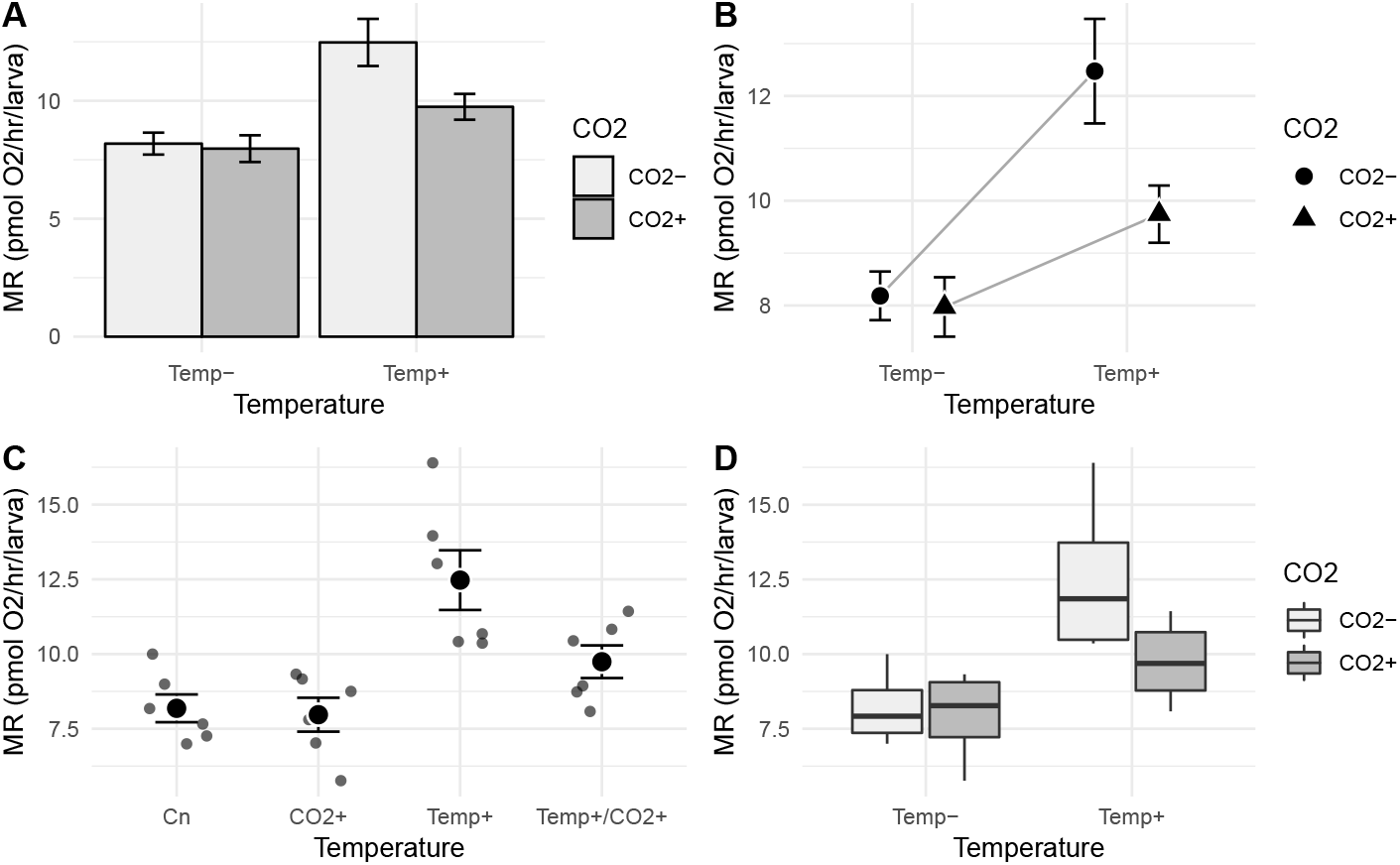
Four alternatives for communicating results of an experiment. (A) bar plot, (B) interaction plot, (C) dot plot, (D) box plot. In all four plot types, the treatment effects are inferred by mentally computing the distance between treatment means (or median in case of the box plot). The interaction effect is inferred in (C) by comparing obliqueness of the lines. The uncertainty in treatment effects and especially the interaction effect cannot be inferred from any of the plots A-D.

### The Harrell plot

The Harrell plot addresses all three recommended practices by combining a forest plot of treatment effects, a box plot, and a jittered dot plot, into a single plot (Figure 1). A Harrell plot has two unconventional features (at least in the ecology, physiology, and cell biology literature) that might confuse a reader at first glance: the plot has two “Y” or response axes, because it is constructed by gluing together two separate plots, and the plot is flipped, so that the response axes are horizontal. The axis labels for the upper plot, which contains the effects, are at the top of the Harrell plot. The axis labels for the lower plot, which contains the raw data, are at the bottom of the Harrell plot.

Modeled effects are illustrated in the upper part of the plot using a dot symbol representing the effect estimate and horizontal bars representing the effect uncertainty. The bars for the urchin data are 95% confidence intervals but these could be credible intervals from a Bayesian analysis. Forest plots of effects with horizontal uncertainty intervals are common in analyses with multiple responses, in meta-analysis, and in the epidemiology literature. The illustrated effects can be the coefficients of the linear model or contrasts between treatment combinations. If contrasts, these can be comparisons with a reference (such as a control) or pairwise comparisons. In the original plot by Harrell that motivated this plot type, the scale of the response axis was the same in the upper and lower plots, so the coordinates in the upper part are a simple, rigid shift of the coordinates in the lower part (Harrell 2014). By contrast, a Harrell plot allows for more flexibility of the scaling of the coordinate system in the upper part, which is either a feature, or a bug, depending on one’s underlying philosophy.

The raw data are shown in the lower part of the plot using jittered dots, clustered by group. The distribution of data in each group is also shown in the lower part of the plot using a box plot. The precise tool to show the data and distributions is flexible but jittered dots and box plot reflect the best practice for much of experimental biology. While some advocate the use of an error bar, the box plot is more informative than an interval based on the sample standard deviation (including the sample confidence interval). And, if we are specifically interested in the effects, a standard error of the mean, or a 95% confidence interval of the mean, is not the uncertainty that we want to communicate (see *Effect size and uncertainty* above).

In the urchin data example (Figure 1), the illustrated effects in the upper part of the Harrell plot are the coefficients of the linear model

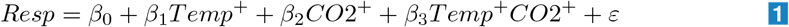

where the dummy variables are coded using treatment (or dummy) coding, which is the default in R but not in many other statistical software packages. Dummy coding is especially useful for this experiment where there is an obvious control or reference level (*Temp^−^/CO*2^−^). In this model, the intercept (*β*_0_) is the mean of this control group. The *Temp*^+^ coefficient (*β*_1_) is the difference in means between the (*Temp*^+^*/CO*2^−^) group and the control group. It is the estimated effect of the added temperature but not added CO2. Similarly, the *CO*2^+^ coefficient (*β*_2_) is the difference in means between the (*Temp^−^/CO*2^+^) group and the control group. It is the estimated effect of the added CO2 but not added temperature. And finally, the interaction coefficient (*β*_3_) is the difference between the mean of the (*Temp*^+^*/CO*2^+^) group and the expected value of this group if the interaction effect were zero, which is *b*_0_ + *b*_1_ + *b*_2_. It’s what’s left to be accounted after adding temperature and CO2.

### Harrell plots vs. common alternatives

Figures 2 A and B illustrate two kinds of mean-and-error plot. The mean response of a group is represented either by the height of the bar (A) or a point symbol (B). The error bar most commonly represents one standard error of the mean (shown here); a confidence interval is less common. Occasionally, the error bar represents one sample standard deviation. Bar-and-error plots are ubiquitous throughout biology. Point-and-error plots like that in (B) are more common in ecology and animal physiology than in cell biology and are sometimes called “interaction plots” because the magnitude of the interaction is inferred by the obliqueness of the lines connecting means. Perhaps because of the ubiquity of bar plots in cell biology (including the pages of Science, Nature, and Cell), most criticism of mean-and-error plots focusses on bar plots, which are pejoratively called “dynamite”, “plunger”, or “antenna” plots.

Three, related criticisms of mean-and-error plots are (Drummond and Vowler 2011; Weissgerber et al. 2015; Rousselet, Foxe, and Bolam 2016), first, they do not show the data, which is important because multiple distributions can produce the same mean and error. Second, mean-and-error plots typically fail to reflect the analyzed model. For example, almost all mean-and-SE plots illustrate standard errors computed independently in each group instead of pooled standard errors resulting from the model. Or, for clustered data (repeated measures, blocked designs) mean-and-error plots give no indication of this lack of independence. And, third, error bars based on the sample standard deviation or standard error of the mean are not easily interpretable, and suggest a false interpretation, if the underlying data are not approximately normal.

Figures 2 C and D illustrate two recommended alternatives to the plots in (A) and (B). (C) is a jittered dot plot (or strip chart) of the raw values within each treatment combination and the treatment means and standard errors. A jittered dot plot is recommended when the treatment combinations have small replicate size, because it “shows the data”. (D) is a box plot, a venerable method developed for exploratory data analysis. A box plot is often recommended when replicate size is large because it summarizes the distribution, including unusual values (or even all values if the raw data are superimposed). Box plots and especially jittered dot plots are becoming increasingly common in cell biology, probably in direct response to the criticisms of bar plots.

The criticisms of mean-and-error plots and even these two solutions do not address the elephant in the room – all four plot types only indirectly communicate what we often want to directly communicate: the effects of the experimental treatments and the uncertainty in these effects. In any of these alternative plots, effects have to be mentally reconstructed by qualitatively comparing the difference in the response between two groups. This is relatively easy (at least at a course level of precision) if there are few groups and if the differences are large relative to the scale of the response axis. In bar plots especially, differences can often be very small relative to the height of the bar and figure. Regardless, effect uncertainty is much harder to mentally reconstruct, because the proper interval is a function of a standard error that is itself a function of the distribution of error variance in multiple groups. Because the approximate end-points of a confidence interval of a difference is time-consuming to mentally construct, mean-and-error plots encourage focus on the presence/absence of an effect (and it’s direction) instead of the magnitude of the effect, including the magnitude of the ends of the confidence interval.

## How Harrell plots improve inference

### Interaction effects

The difficulty with reconstructing effects from the four plot types is magnified when considering an interaction effect. A common practice throughout biology (encouraged by textbooks and especially the applied literature) is to use the *p*-value of an interaction term in an ANOVA table in a decision rule to choose one of two conclusions: 1) the interaction *p* is significant and main effects are not interpreted but, perhaps, conditional (simple) effects are, or 2) the interaction *p* is not significant (and often mistakenly reported as “no effect”) and the main effects are interpreted from the same table. Occasionally (and more common in ecology), the non-significant interaction *p*-value is the signal to refit the model without the interaction term before interpreting the main effects.

Any of the four alternative plots (bar, interaction, dot, box) encourage this dichotomization strategy because none directly communicates the interaction effect and its uncertainty. By contrast, a Harrell plot encourages best practices. The 95% CI of the interaction effect of the urchin data includes large negative values at one end but only trivially small positive values at the opposite end. These data, then, are not consistent with a moderate or large positive interaction, but are consistent with large negative values. Indeed, the maximum likelihood estimate of the interaction effect is moderately large – over 1/2 the magnitude of the estimated *Temp* effect. Given the results presented in the Harrell plot, it just doesn’t seem very sound to follow the *p*-value decision rule outlined above and conclude that there is “no interaction effect” and that temperature and CO_2_ act additively on urchin metabolism.

This lack of soundness is highlighted by comparing the Harrell plot in Figure 3B to that in Figure 3A. The data used for Figure 3B are the original, simulated data with the residual standard deviation contracted from 1.55 to 1.4 pmol/hr/larva (10%) and interaction effect increased from −2.6 to −2.8 (7.7%). The consequence of this shift is the interaction *p* = 0.037 and the CI of the interaction effect not covering zero. If we used the interaction *p* decision rule, these two results would be interpreted quite differently! But if we were to use the Harrell plots, we would interpret these the same: the interaction effect is moderately negative and both large, negative values and very small values are consistent with the data. In other words, if we had an *a priori* model that predicted no interaction (or a strong interaction), then the results are too ambiguous. While the best estimate is a moderately large, negative interaction, a trivially small interaction is consistent with the data.

**Figure 3:**
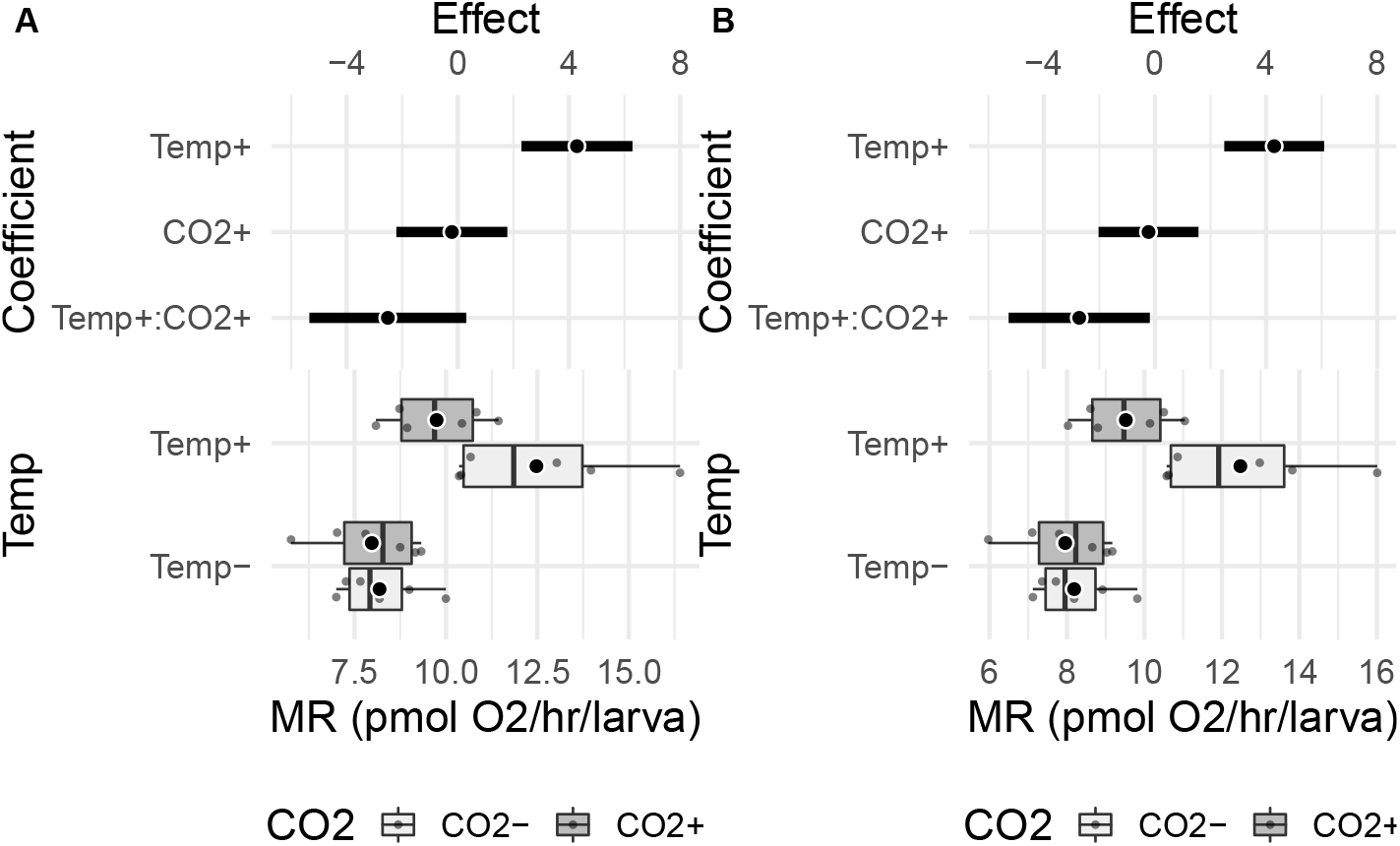
A tale of two plots. The Harrell plot in (A) is the original, simulated data, with interaction *p* = 0.077. The data in the Harrell plot in (B) is a slight modification of that in (A), with the consequence that the interaction *p* = 0.037.

### Harrell plots encourage focusing on effect magnitude and uncertainty instead of levels of effects

The epidemic of *p*-values that invariably results from reporting ANOVA tables has spread to plots, where asterisks indicating “levels of significance” and letters classifying groups into statistically different bins are now common. Figure 4A is a bar plot with 1 SE bar for the results of a *Drosophila* wind-tunnel selection experiment. The unpublished data are the maximum burst speed of *Drosophila melanogaster* individuals from two lines (AA) that have undergone selection in a compartmentalized wind tunnel and two control lines (CN) (Marden, Wolf, and Weber 1997). Maximum burst speed was measured on individual flies that were stimulated to take-off and fly against a wind of known speed in a wind tunnel. The asterisks, which “levels” of significance for tests of specific contrasts, draw the researcher and reader into comparing means using the heights of the bars. The asterisks are helpful for this because the standard error bars are not especially helpful given the computations necessary to go from the illustrated standard error of *means* to the relevant standard error of *differences*. But the asterisks encourage the researcher/reader into focussing on *p*-values of inferential tests. Even more egregiously, the asterisks encourage the researcher/reader to misinterpret the number of asterisks as an effect size.

**Figure 4:**
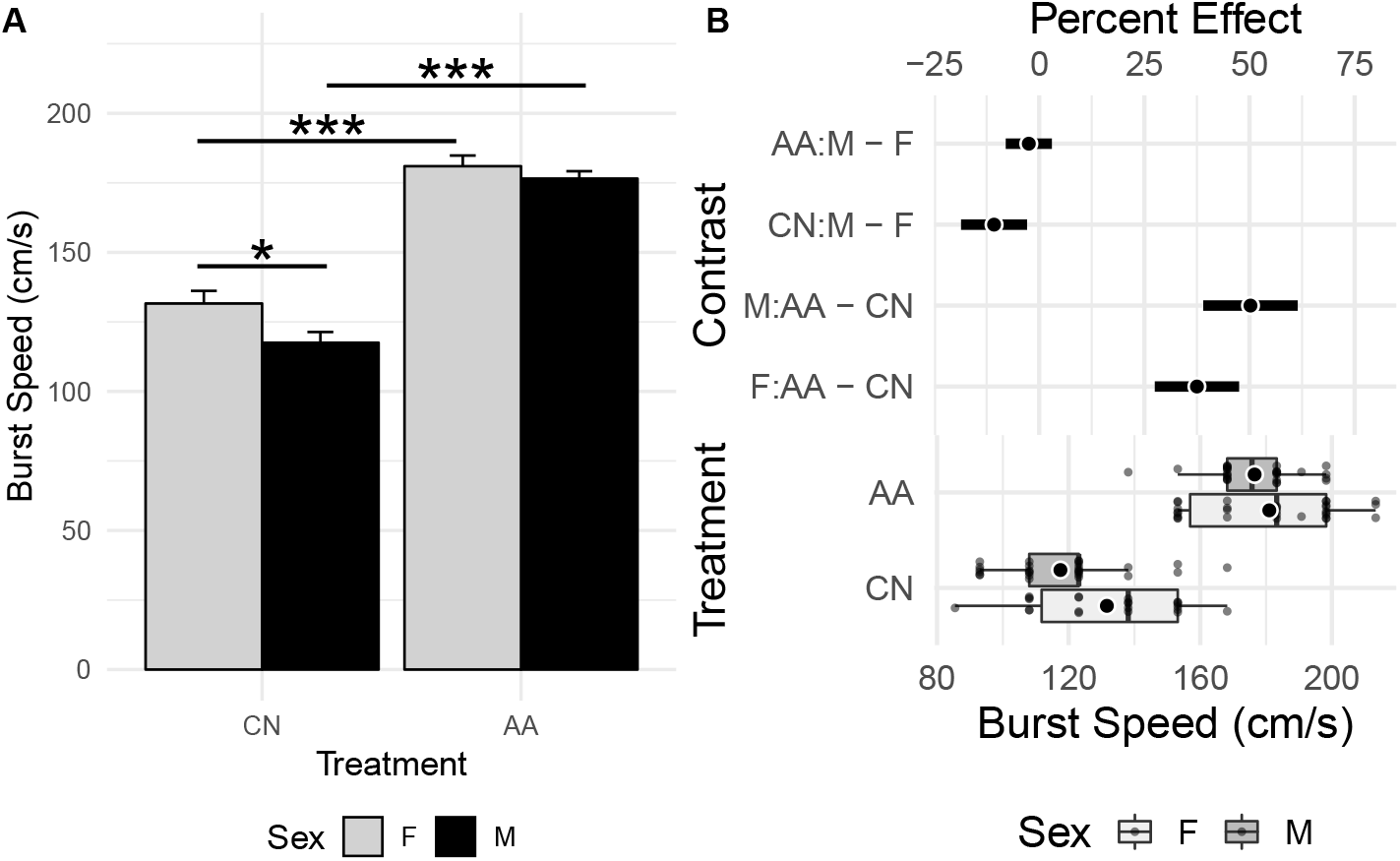
Levels of significance vs effects and uncertainty. (A) Bar plot of fly data. Error bars are 1 SEM. Asterisks indicate level of significance of inference test. (B) Harrell plot of fly linear model results. Conditional (simple) effects are shown in the upper part. Bars are 95% confidence intervals. The effects in the upper part are contrasts and the – in the labels is a minus sign and not a hyphen. Key: control lines (CN), selected lines (AA), male (M), female (F).

A Harrell plot (Figure 4B), by contrast, nudges researchers/readers to focus on modeled effect size and uncertainty instead of classification of results into “significant” and “non-significant” bins (or “highly significant” bins). Here, I’ve scaled the contrasts as percent differences 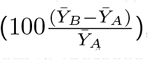, because I haven’t given much thought to the biological consequences of a 50 cm/s increase in burst speed. Thinking about effect size as a percent difference has several downsides, including, 1) while effect size scaled as percents give us some sense of importance, percents discourage the hard work of understanding the biological consequences of absolute effect size, and 2) the standard error of the effect scaled as a percent is not simply 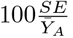 (and is larger than this value). One can use the 95% confidence limits to mentally bin comparisons into significant and non-significant bins, but the plot itself does not encourage it. Instead, the plot encourages a researcher/reader to consider how the range of values in a CI support, or fail to support, different interpretations of the results. For example, the effects of the long term selection experiment is much larger than the effects of sex on burst speed. And, a model of males having smaller burst speed relative to females in both wind-tunnel selected (AA) and control (CN) flies is consistant with the data.

### Harrell plots show modeled effects

In Figure 5, I have redrawn Figure 3 from Kardol, Spitzer, et al. (2016b), which shows treatment means and standard errors for moss biomass in response to different combinations of community complexity and precipitation. The original data are available from the Dryad Digital Repository (Kardol, Spitzer, et al. 2016a). Different letters classify the treatment levels into statistically distinct bins using significant *p*-values for tests of marginal means. In general, the magnitude and uncertainty in contrasts of marginal means (or “main” effects) in factorial designs are difficult to infer from any kind of mean-and-error plots of cell means, such as Figure 5 or any of those in Figure 2, because this requires mentally averaging over multiple levels. Or at least the ease of this decreases rapidly with number of treatment levels and heterogeneity of conditional effects.

**Figure 5:**
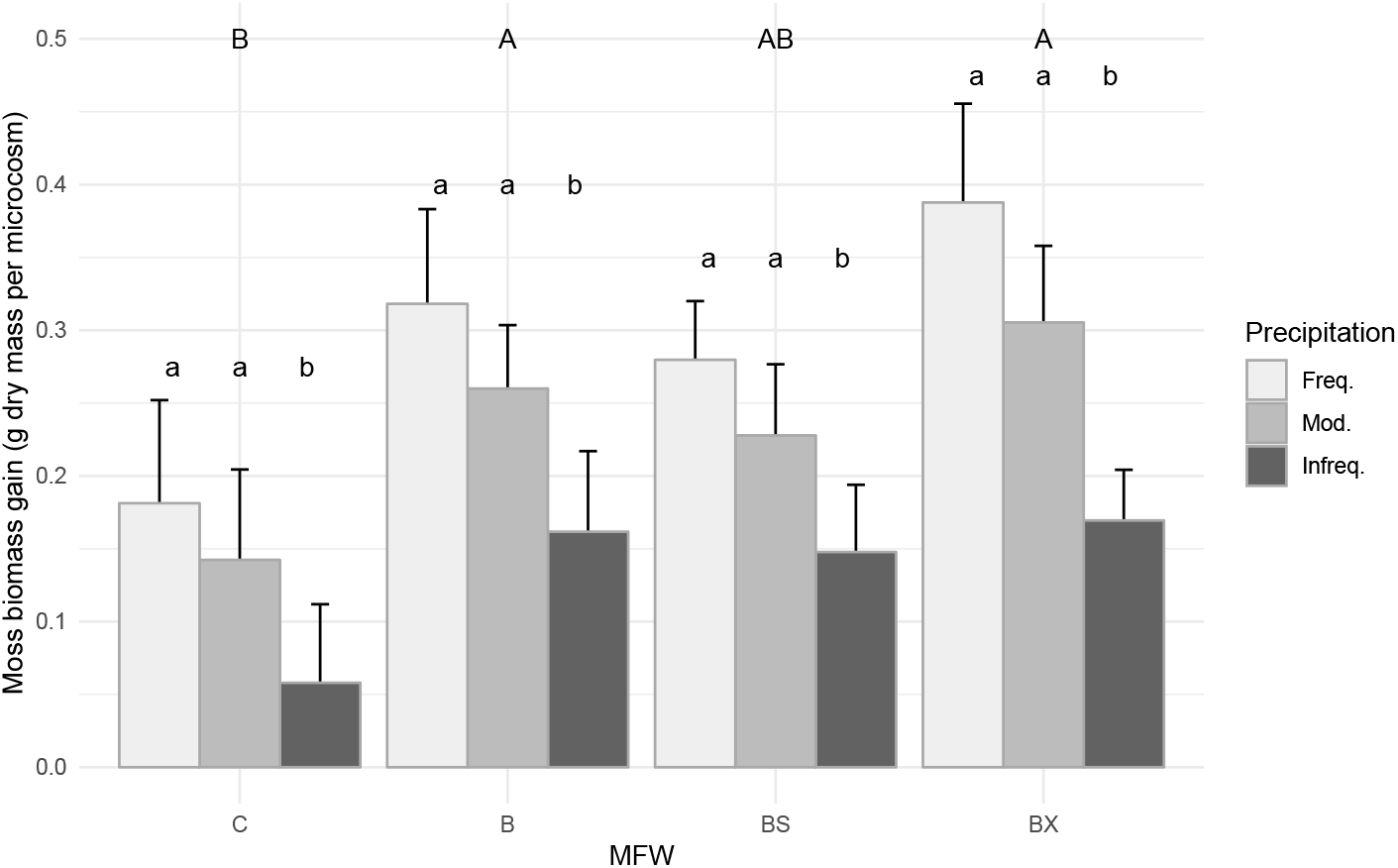
Bar plot of moss data. Error bars are 1 SEM. Letters indicate statistically significant tests of marginal means pooled over levels of other factor

By contrast, a Harrell plot of the data and linear model (Figure 6 directly shows the modeled differences among marginal means with a measure of the uncertainty of these estimates (95% confidence intervals). Certainly, one can use the 95% confidence limits to mentally classify comparisons into significant and non-significant bins, but the plot itself does not encourage it. Instead, the plot encourages a researcher/reader to consider how the range of values in a CI support, or fail to support, different interpretations of the results. For example, trivial to moderate (50%) reductions in biomass as rain decreases is consistent with the *Moderate – Frequent* contrast CI, but trivial to moderate increases in biomass as rain decreases are not consistent with the CI. Means with standard errors supplemented with letters or asterisks do not convey this information.

**Figure 6:**
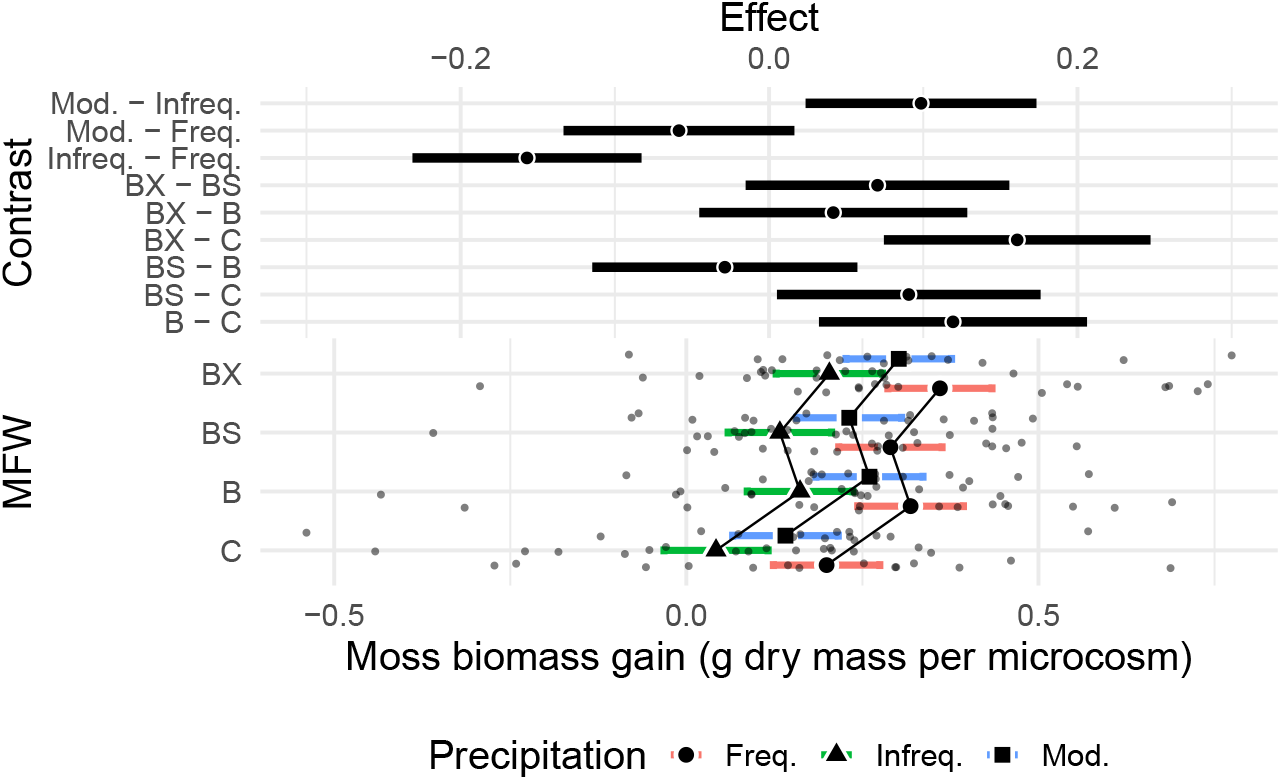
Harrell plot of moss linear model results. The effects in the upper part are differences between marginal means. The treatment means in the lower part are conditional on additive effects only. Error bars of marginal effects and modeled means are 95% confidence intervals

One needs only the upper, forest-plot component of the Harrell Plot to communicate treatment effects and their uncertainty but the lower, treatment summary plot also communicates important information. In Figure 6, the illustrated means and confidence intervals are are conditioned only on the additive effects, which is why the lines connecting the means are parallel. The magnitudes of the cell or modeled means are often meaningful to knowledgeable readers (for example, I know the range of fast start acceleration in fish so if I see plots with data outside of this range, this will immediately raise a red flag about the analysis). Sometimes, the treatment means are directly relevant for interpretation or decision making. Other times, the dots or box plots simply give us a since of what the data are. Here, for example, the dot/box plot indicates that the response (moss biomass gain) can take negative values while the bar plot in Figure 5 fails to indicate this. Given only the bar plot, a reader might mistakenly infer that the response is a count, or some other variable that can only take positive values.

### Conclusion: Harrell plots show the model

A spate of articles, editorials, and blog posts have criticized bar plots because they fail to show the underlying data and have advocated replacing bar plots with dot plots or box plots. What all these plots fail to show is the modeled effects and uncertainty, which is the most important result of an experiment. A Harrell plot shows not only “the data” but also “the model”, or at least the model results. A Harrell plot encourages best practices by nudging researchers and readers to focus on effect size and to interpret the whole confidence interval and not just assessing if the interval includes or excludes zero (or some non-nill null). While a Harrell plot can lead a researcher to good science, it can’t make them think. For that, researchers will need to do the hard work of working out the ecological, physiological, and health consequences of effect size.

## References

Amrhein, Valentin, Fränzi Korner-Nievergelt, and Tobias Roth. 2017. “The Earth Is Flat (P *>* 0.05): Significance Thresholds and the Crisis of Unreplicable Research.” PeerJ 5 (July). doi:10.7717/peerj.3544.

Batterham, Alan M., and William G. Hopkins. 2006. “Making Meaningful Inferences About Magnitudes.” International Journal of Sports Physiology and Performance 1 (1):50–57.

Curran-Everett, Douglas, Sue Taylor, and Karen Kafadar. 1998. “Fundamental Concepts in Statistics: Elucidation and Illustration.” Journal of Applied Physiology 85 (3):775–86.

Drummond, G. B., and S. L. Vowler. 2011. “Show the Data, Don’t Conceal Them.” The Journal of Physiology 589 (8):1861–3. doi:10.1113/jphysiol.2011.205062.

Gelman, Andrew. 2013. “P Values and Statistical Practice:” Epidemiology 24 (1):69–72. doi:10.1097/EDE.0b013e31827886f7.

Gelman, Andrew, and John Carlin. 2014. “Beyond Power Calculations: Assessing Type S (Sign) and Type M (Magnitude) Errors.” Perspectives on Psychological Science 9 (6):641–51. doi:10.1177/1745691614551642.

Greenland, Sander, and Charles Poole. 2013. “Living with P Values.” Epidemiology 24 (1):62–68. doi:10.1097/EDE.0b013e3182785741.

Harrell, Frank E. 2014. “Principles of Graph Construction.”

Johnson, Douglas H. 1999. “The Insignificance of Statistical Significance Testing.” The Journal of Wildlife Management, 763–72.

Kaelin, Jr William G. 2017. “Publish Houses of Brick, Not Mansions of Straw.” Nature News 545 (7655):387. doi:10.1038/545387a.

Kardol, Paul, C.M. Spitzer, Gundale, M. J., Nilsson, M, and Wardle, D. A. 2016a. “Data from: Trophic Cascades in the Bryosphere: The Impact of Global Change Factors on Top-down Control of Cyanobacterial N2-Fixation.” Dryad Digital Repository. doi:10.5061/dryad.66d5f.

Kardol, Paul, Clydecia M. Spitzer, Michael J. Gundale, Marie-Charlotte Nilsson, and David A. Wardle. 2016b. “Trophic Cascades in the Bryosphere: The Impact of Global Change Factors on Top-down Control of Cyanobacterial N _2_-Fixation.” Edited by Mark Gessner. Ecology Letters 19 (8):967–76. doi:10.1111/ele.12635.

Krzywinski, Martin, and Naomi Altman. 2014. “Points of Significance: Visualizing Samples with Box Plots.” Nature Methods 11 (2):119–20.

Marden, J. H., M. R. Wolf, and K. E. Weber. 1997. “Aerial Performance of Drosophila Melanogaster from Populations Selected for Upwind Flight Ability.” Journal of Experimental Biology 200 (21):2747–55.

Munafò, Marcus R., and George Davey Smith. 2018. “Robust Research Needs Many Lines of Evidence.” News. Nature. http://www.nature.com/articles/d41586-018-01023-3. doi:10.1038/d41586-018-01023-3.

Nakagawa, S, and I C Cuthill. 2007. “Effect Size, Confidence Interval and Statistical Significance: A Practical Guide for Biologists.” Biological Reviews, January, 591–605. doi:10.1111/j.1469-185x.2007.00027.x.

Padilla-Gamiño, Jacqueline L., Morgan W. Kelly, Tyler G. Evans, and Gretchen E. Hofmann. 2013. “Temperature and Co2 Additively Regulate Physiology, Morphology and Genomic Responses of Larval Sea Urchins, Strongylocentrotus Purpuratus.” Proceedings of the Royal Society of London B: Biological Sciences 280 (1759):20130155. doi:10.1098/rspb.2013.0155.

Rousselet, Guillaume A., John J. Foxe, and J. Paul Bolam. 2016. “A Few Simple Steps to Improve the Description of Group Results in Neuroscience.” European Journal of Neuroscience 44 (9):2647–51. doi:10.1111/ejn.13400.

Spitzer, Michaela, Jan Wildenhain, Juri Rappsilber, and Mike Tyers. 2014. “BoxPlotR: A Web Tool for Generation of Box Plots.” Nature Methods 11 (2):121–22.

Tukey, John W. 1991. “The Philosophy of Multiple Comparisons.” Statistical Science 6 (1):100–116.

Vaux, David L. 2012. “Research Methods: Know When Your Numbers Are Significant.” Nature 492 (7428):180–81. doi:10.1038/492180a.

Weissgerber, Tracey L, Vesna D Garovic, Stacey J Winham, Natasa M Milic, and Eric M Prager. 2016. “Transparent Reporting for Reproducible Science.” Journal of Neuroscience Research 94 (10):859–64. doi:10.1002/jnr.23785.

Weissgerber, Tracey L., Natasa M. Milic, Stacey J. Winham, and Vesna D. Garovic. 2015. “Beyond Bar and Line Graphs: Time for a New Data Presentation Paradigm.” PLOS Biology 13 (4):e1002128. doi:10.1371/journal.pbio.1002128.

Yoccoz, Nigel G. 1991. “Use, Overuse, and Misuse of Significance Tests in Evolutionary Biology and Ecology.” Bulletin of the Ecological Society of America 72 (2):106–11.

